# The cGAS-STING pathway is an *in vivo* modifier of genomic instability syndromes

**DOI:** 10.1101/2024.10.16.618655

**Authors:** Marva Bergman, Uri Goshtchevsky, Tehila Atlan, Gwendoline Astre, Ryan Halabi, Hosniyah El, Eitan Moses, Aaron J.J. Lemus, Bérénice A. Benayoun, Yehuda Tzfati, Ido Ben-Ami, Itamar Harel

**Affiliations:** Department of Genetics, the Silberman Institute, the Hebrew University of Jerusalem, Givat Ram, Jerusalem, 91904, Israel; Department of Obstetrics & Gynecology, Shaare Zedek Medical Center and Faculty of Medicine, The Hebrew University of Jerusalem, Jerusalem, Israel; Leonard Davis School of Gerontology, University of Southern California, Los Angeles, CA 90089, USA; Molecular and Computational Biology Department, USC Dornsife College of Letters, Arts, and Sciences, Los Angeles, CA 90089, USA

## Abstract

Mutations in genes involved in DNA damage repair (DDR) often lead to premature aging syndromes. While recent evidence suggests that inflammation, alongside mutation accumulation and cell death, may drive disease phenotypes, its precise contribution to *in vivo* pathophysiology remains unclear. Here, by modeling Ataxia Telangiectasia (A-T) and Bloom Syndrome in the African turquoise killifish (*N. furzeri*), we replicate key phenotypes of DDR syndromes, including infertility, cytoplasmic DNA fragments, and reduced lifespan. The link between DDR defects and inflammation is attributed to the activation of the cGAS-STING pathway and interferon signaling by cytoplasmic DNA. Accordingly, mutating cGAS partially rescues germline defects and senescence in A-T fish. Double mutants also display reversal of telomere abnormalities and suppression of transposable elements, underscoring cGAS’s non-canonical role as a DDR inhibitor. Our findings emphasize the role of interferon signaling in A-T pathology and identify the cGAS-STING pathway as a potential therapeutic target for genomic instability syndromes.

## Introduction

Genomic instability is a central player in aging and age-related diseases^1–4^. This aging hallmark encompasses various naturally occurring DNA mutations, including base substitutions, copy number variations, chromosomal aberrations, and deletions. Subsequently, the DNA damage response (DDR) coordinates the intricate repair process through a network of sensors, transducers, and effectors^5^. Accordingly, mutations that impair the DDR, particularly those involved in the response to double-strand breaks (DSBs), underlie various human progeroid syndromes. For example, mutations in *blm, wrn,* and *atm* genes can give rise to Bloom syndrome (BS), Werner syndrome (WRN), and Ataxia telangiectasia (A-T)^6–8^, respectively.

Different DDR-related syndromes can display overlapping and distinct pathologies^1–3^, including predominantly cancer predisposition (e.g., BRCA^7,9^) or segmental accelerated aging (e.g., A-T^10^). This heterogeneity is thought to be mediated by the nature of each lesion, which can be mostly mutagenic (e.g., oxidative damage) or cytotoxic (e.g., DSB), as well as by the cellular state (e.g. proliferative or differentiated). Recent evidence suggests that DNA-damage-induced inflammation^11–15^ may drive disease progression, in addition to mutation accumulation and cell death. However, the precise contribution of each process to disease phenotypes remains unclear. Understanding the interaction between the accumulation of cellular damage and its systemic effects through the immune system may offer a theoretical framework for the role of genome maintenance in aging and disease^1,2,6,7,16^.

In addition to mutation accumulation and cell death, DNA damage can exert a variety of cellular and physiological effects. For example, polymerase stalling following DNA damage can reduce transcript expression, specifically in longer genes^17^. Another example is the increase in the cyclic GMP– AMP synthase (cGAS)–stimulator of interferon genes (STING) pathway following DNA damage^11–15^. Specifically, unrepaired DNA fragments that enter the cytoplasm are perceived as an invading pathogen, and trigger an immune response through dedicated sensors (e.g., cGAS^11,18^). Subsequently, innate immunity is activated through the type-I interferon (IFN-I) system and proinflammatory cytokines^13^.

Activation of the cGAS–STING pathway contributes to many age-related changes^4,16^, including chronic age-related inflammation, also known as ‘inflammaging’^19^, which can exacerbate various age-related pathologies^11–15,18,20,21^. While this pathway is required for normal health^22,23^, inhibiting the cGAS-STING pathway was proposed to have a beneficial effect in inflammatory diseases^15^, senescence^24–26^, neurodegeneration^20^, and genomic instability in cultured organoids^27^. However, whether modulation of the cGAS-STING pathway can rescue specific pathologies linked with genomic instability *in vivo* remains underexplored.

Many murine models have been successfully developed to investigate genomic instability syndromes, including BS and A-T^9^. However, in some cases, the phenotypic spectrum is incomplete. For example, because the telomeres of laboratory mice are unusually long, the mouse model for WRN displays clinical features only when bred on a telomerase-deficient background^28^. Similarly, as *blm* mutants in mice are embryonic lethal^29^, conditional inactivation is required^30^. Zebrafish DDR models have recently emerged as exciting alternatives^31–34^, yet, the relatively long lifespan of both vertebrate models remains an experimental challenge^35^.

Here, we leverage the turquoise killifish (*Nothobranchius furzeri*) as an experimental platform to identify functional modifiers of genomic instability. The killifish has recently emerged as a promising genetic model for aging, owing to a naturally compressed lifespan (∼6-10-fold shorter than mice and zebrafish, respectively^35^), and the availability of state-of-the-art genome editing tools^36–43^. These features have enabled the identification of novel vertebrate longevity mechanisms (through the AMP/AMPK pathway or via germline manipulations^41,42,44^), and the rapid modeling of human age-related syndromes (e.g. telomere syndrome^36^).

We first generate two genetic killifish models for DSB-related progeroid syndromes, the Bloom syndrome (BS) and Ataxia telangiectasia (A-T). Our models faithfully recapitulate key human pathologies, including infertility, reduced lifespan, modified immunity, and various cellular phenotypes. To investigate the contribution of the cGAS-STING pathway to disease progression, we genetically inactivate the cytosolic DNA sensor cGAS. Interestingly, *atm;cgas* double mutants exhibit rescued disease phenotypes, including reduced inflammation and senescence, and restored germline development. We also show that cGAS inhibition itself can negatively affect genomic stability. Our findings propose the cGAS–STING pathway as a promising *in-vivo* target for treating DDR syndromes, and encourage further investigations concerning its safety.

## Results

### Generating genetic models for DDR in killifish

To functionally explore the role of DSBs, we generated two genetic models for DDR-related syndromes. Using our recent CRISPR protocols^45,46^ we edited the *atm* and *blm* genes, generating frameshift deletions that are predicted to be loss-of-function alleles (*atm^Δ4^* and *blm^Δ11^*, **Figure S1a**). We then outcrossed heterozygous fish for several generations to reduce the burden of possible off-target mutations. Mating heterozygous pairs followed the expected Mendelian ratios concerning the fish genotypes (p = 0.4 for *blm*, and p=0.09 for *atm*, **Figure S1b**), consistent with the absence of a significant effect on embryonic development. We next produced homozygous mutants, for both alleles and set to explore physiological and cellular phenotypes characteristic of the human syndromes.

### Hepatic transcriptional signatures of *blm* mutants suggest conserved disease phenotypes

Bloom Syndrome patients suffer from early onset of age-related diseases, including reduced fertility, cancer predisposition, and immune and metabolic abnormalities. Therefore, we initially characterized the hepatic transcriptome of *blm^Δ11/Δ11^*fish. Principal component analysis (PCA) suggested that the transcriptomes of WT and *blm* livers are separated (**Figure 1a**), further supported by 74 downregulated and 541 upregulated genes (FDR < 0.01, **Table S1**).

**Figure 1:**
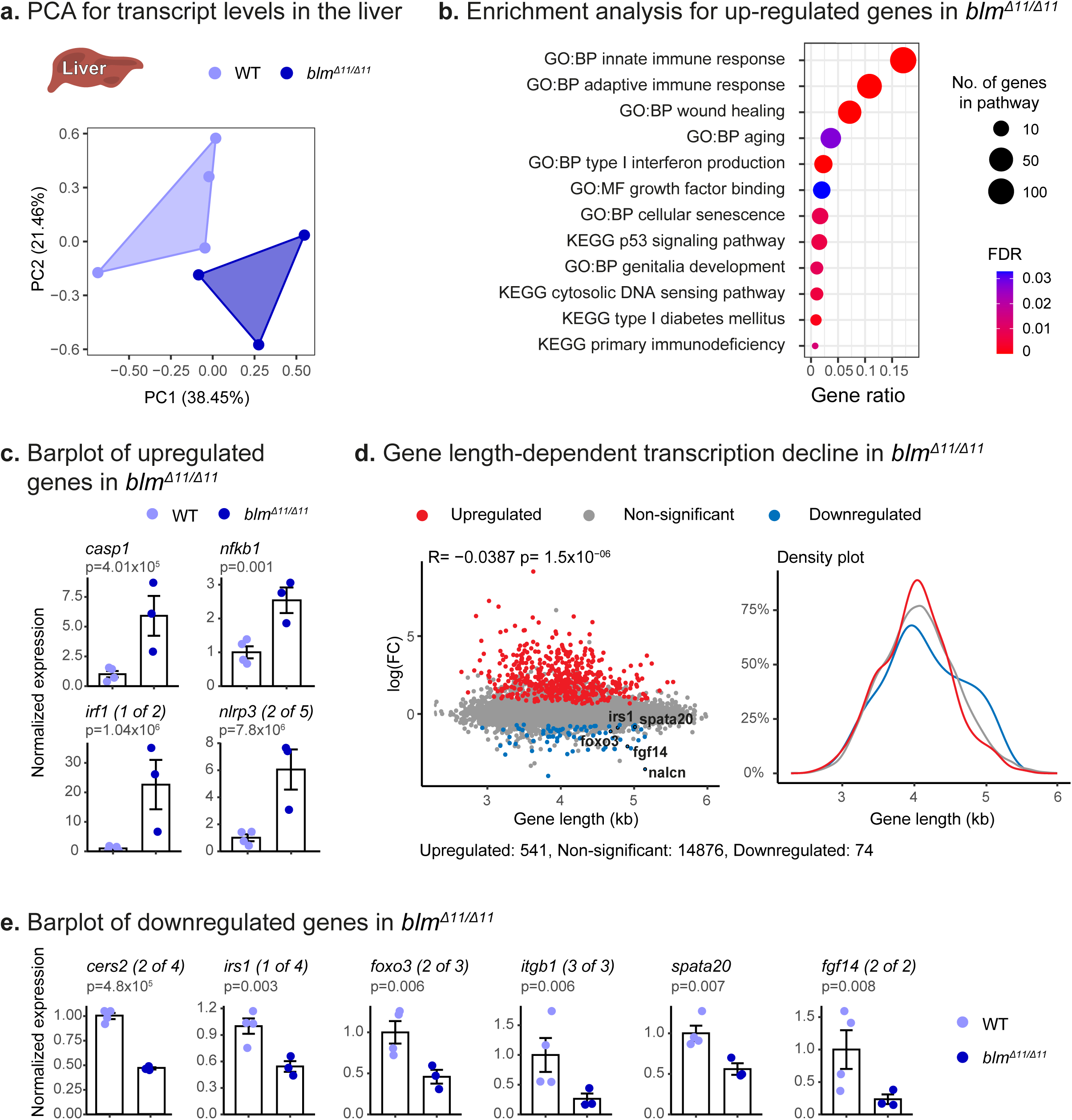
Transcriptional characterization of the liver in the killifish model for Bloom Syndrome. **(a)** PCA of hepatic transcript levels. *n* = 3-4 samples per condition, with each symbol representing an individual fish. **(b)** Functional enrichment analysis (GO) for upregulated genes between WT and *blm^Δ11/Δ11^* mutant fish (*p* < 0.01). GO enrichments were identified at FDR < 5%. **(c)** Relative expression of driver genes from the pathways in (b). For each gene, the expression level is normalized to WT. Significance was called as part of the edgeR pipeline using the classic mode. **(d)** Left: Scatter plot of gene length versus log-fold change between WT and *blm^Δ11/Δ11^*. *R*-value and *p*-value are indicated. Right: a density plot of gene length that separates significant up-or down-regulated genes. **(e)** Relative expression of selected long genes from (d). For each gene, the expression level is normalized to WT. Significance was determined using the edgeR pipeline in classic mode.

We then performed GO enrichment analysis on the significantly upregulated genes, implicating the involvement of various pathways associated with typical BS disease phenotypes. These pathways highlighted deregulated immune functions (e.g. interferon type I, immunodeficiency), growth and reproduction, cellular senescence, and metabolic disorders (e.g. diabetes, **Figure 1b**). Selected upregulated driver genes that are part of the inflammatory and interferon response are also indicated (**Figure 1c**).

Gene length-dependent transcription decline was recently suggested to be linked to polymerase stalling following DNA damage^17,47,48^. Accordingly, further examination of significantly downregulated genes suggested they were longer than expected (**Figure 1d, e**, and **Table S1**). Importantly, these genes are associated with disease-related functions, including aging^49^ (*Forkhead Box O3,* or *foxo3*), type II diabetes^50^ (*Insulin Receptor Substrate 1,* or *irs1*), and spermatogenesis (*Spermatogenesis Associated 20,* or *spata20*).

Together, these transcriptomic findings imply that classical BS disease phenotypes are conserved in the killifish *blm* mutants. Furthermore, functions regulated by longer genes could indirectly contribute to the disease progression. Next, we explored selected disease phenotypes that are characteristic of DDR syndromes.

### Killifish models for DDR display reproductive defects

BS and A-T patients suffer from disease phenotypes at different degrees of severity, including infertility and growth retardation^51–54^. Characterizing the homozygous *atm^Δ4/Δ4^* and *blm^Δ11/Δ11^*killifish mutants revealed that these fish displayed a modest effect on growth, particularly in *blm^Δ11/Δ11^* fish (**Figure S2a**). To evaluate killifish reproductive phenotypes^41^, we stained gonadal tissue sections with Hematoxylin and Eosin (H&E). Our findings indicate that while *blm^Δ11/Δ11^* fish experienced a milder phenotype, with a reduction in the number of mature eggs and sperm compared to WT fish, mature oocytes and sperm could not be detected at all in *atm^Δ4/Δ4^* fish (**Figures 2a, b**). Our findings were further confirmed by mating homozygous pairs, indicating that *blm^Δ11/Δ11^* pairs displayed reduced fertility, while *atm^Δ4/Δ4^* fish were completely infertile (**Figure S2b**).

**Figure 2:**
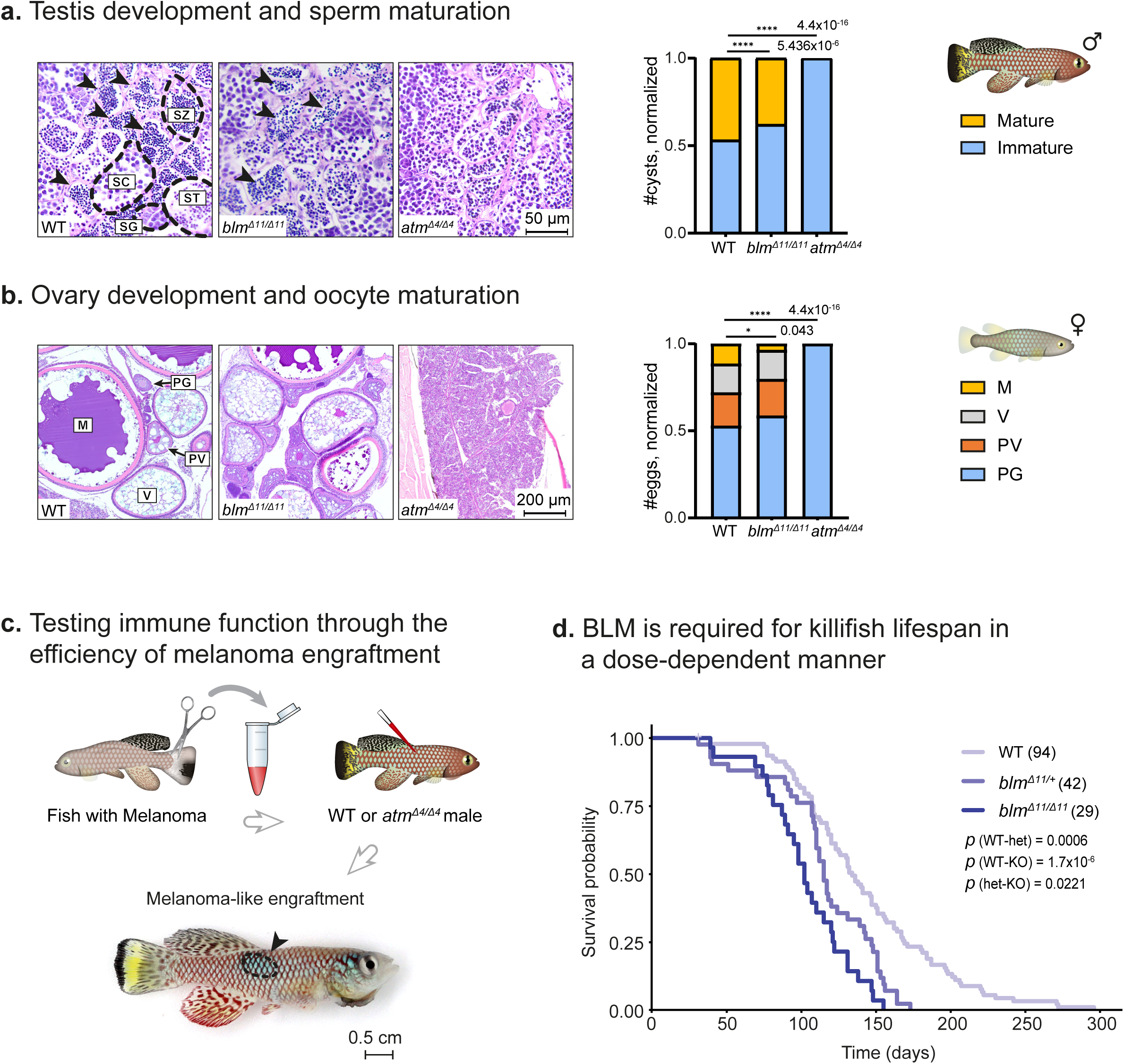
Killifish DDR mutants replicate physiological features of genomic instability syndromes. **(a)** Left: Histological sections from three-month-old males. *n* ≥ 4 individuals per genotype. Scale bar: 50 µm. Sperm developmental stages (according to^41^): SG (spermatogonia), SC (spermatocytes), ST (spermatids), SZ (spermatozoa, also marked by black arrows). Right: Quantification of sperm maturation stages, presented as a proportion of mature and immature sperm. Significance was assessed using a χ² test with the WT as the expected model, and FDR correction. **(b)** Left: Representative histological sections showing ovaries from three-month-old females of the indicated genotypes. *n* ≥ 5 individuals per genotype. Scale bar: 200 µm. Oocyte development stages (according to^41^): PG (primary growth), PV (pre-vitellogenic), V (vitellogenic), M (mature). Right: Quantification of oocyte maturation stages for each genotype. Data are presented as the proportion of each oocyte developmental stage. Significance was assessed using a χ² test with the WT as the expected model, with FDR correction. **(c)** Experimental design for engrafting melanoma-like cells into *atm ^Δ4/Δ4^*mutant recipients. n ≥ 5 biological replicates. Findings were compared to WT and *rag2* mutants in^43^. Scale: 0.5cm **(d)** Lifespan of WT, *blm^Δ11/+^*, and *blm^Δ11/Δ11^* fish, assessed for both sexes together (separate graphs are in S2). *p* values for differential survival in log-rank tests and fish numbers are indicated.

### Cancer engraftment in *atm^Δ4/Δ4^* mutants suggest immunodeficiency

A-T patients are highly predisposed to malignancies^10^, and accordingly, we have recently identified an increased incidence of sporadic age-related cancers in killifish *atm^Δ4/Δ4^* mutants^43^. While cancers are classically associated with mutation accumulation in DDR syndromes, cancer predisposition could also be linked to another feature of the A-T syndrome, immunodeficiency^10^.

As the interplay between immunodeficiency, mutation accumulation, and cancer predisposition is difficult to untangle, we applied a cancer transplantation assay into *atm^Δ4/Δ4^* mutants to directly evaluate the contribution of the immune system (**Figure 2c**). Specifically, we injected a melanoma-like killifish tumor^43^ into *atm^Δ4/Δ4^* mutant fish and evaluated visible tumors 5 weeks post-injection (**Figure 2c**). Of the surviving fish at week 5, 100% of *atm^Δ4/Δ4^*mutants (5/5) exhibited tumor-like expansions around the injection site (compared to only 50% of WT fish^43^).

Interestingly, the high cancer engraftment in *atm^Δ4/Δ4^* recipients is comparable to rates observed in immunodeficient killifish *rag2^null^* recipients, which lack functional B-and T-cells^43^. Thus, suggesting that modified immune functions in A-T patients could play an important role in cancer predisposition.

### DDR is required for killifish lifespan in a dose-dependent manner

Given the central role of DNA damage in aging, we were curious to examine the lifespan of our DDR models. In agreement with recent findings^34^ the lifespan of *blm^Δ11/Δ11^* homozygous mutants was significantly shorter compared to WT controls in both sexes (∼24% decrease in median lifespan, log-rank test, p = 1.7X10^-6^, **Figures 2d**, **S2c**). Similar trends were also observed for *atm^Δ4/Δ4^* mutants when stratified by sex (in females, log-rank test, *p* = 0.034, **Figure S2c**). Interestingly, measuring the lifespan of *blm^Δ11/+^* heterozygous fish revealed that they undergo an intermediate phenotype (∼14% decrease in median lifespan from WT, log-rank test, *p* = 0.0006, **Figure 2d**, **Table S1**). Thus, our data demonstrates that ATM and BLM are required for a killifish lifespan, and BLM in a dose-dependent manner.

### Altered cellular DDR in mutant-derived primary cells

In mammals, the ATM protein plays a key step in the initiation of the DDR through histone H2AX phosphorylation (γ-H2AX). Additional ATM substrates include BRCA1, BLM, and 53BP1^55–57^. Therefore, to explore the cellular DNA damage response we generated primary killifish fibroblast cultures^42,43^ derived from either WT or DDR-mutant fish (**Figure 3a**).

**Figure 3:**
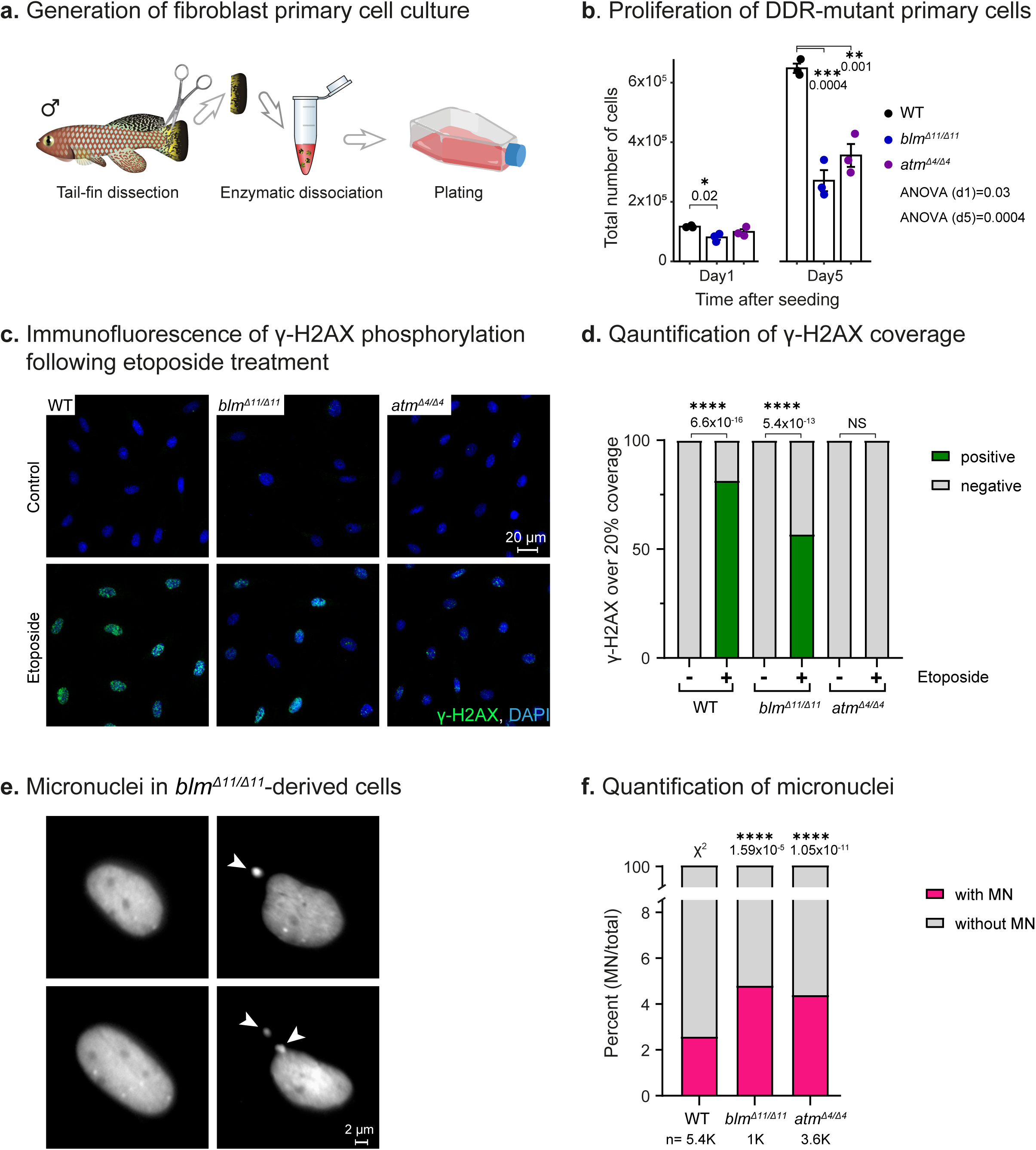
Conserved cellular phenotypes in primary cells derived from DDR mutant fish. **(a)** Schematic illustration of the isolation procedure for primary fibroblasts from the tail fin of individual fish (*n* ≥ 3 for each genotype). **(b)** Quantification of primary fibroblast proliferation from DDR mutants, 1 and 5 days after seeding. Significance was calculated using one-way ANOVA with Tukey post hoc, and *p* values are indicated. Error bars show mean ± SEM. **(c)** Immunofluorescence staining of primary fibroblast cultures, either with or without Etoposide treatment, stained for γ-H2AX and DNA (DAPI). Scale bar: 20 µm. **(d)** Quantification of γ-H2AX nuclear coverage (as a proxy for foci number). Nuclei with over 20% coverage were considered positive. Significance was calculated using Fisher’s exact test and FDR correction. **(e)** Micrographs of DAPI stained *blm^Δ11/Δ11^*-derived primary fibroblasts. Cells with micronuclei are shown with white arrows. **(f)** Quantification of micronuclei. The percentage of micronuclei-positive cells is presented for each genotype. Significance was calculated using χ^2^ proportions test with WT proportions as expected values and FDR correction.

As expected, both *atm^Δ4/Δ4^*-and *blm^Δ11/Δ11^*-derived cells display reduced proliferation compared to WT cells (**Figure 3b**). We next induced DNA damage by etoposide exposure and performed immunofluorescence for γ-H2AX (**Figure 3c**). Our data indicated that, as expected^55–57^, *atm^Δ4/Δ4^*-derived cells fail to phosphorylate H2AX in response to DNA damage, while *blm^Δ11/Δ11^*-derived cells displayed a reduction in γ-H2AX foci (**Figure 3c, d**).

As a result of genomic instability, DNA fragments can be erroneously separated from the primary chromosomal mass, and segregated as distinct structures known as micronuclei^58,59^. Quantifying micronuclei using a nuclear dye (DAPI, **Figures 3e** and **S3a**) revealed an increase in both *atm^Δ4/Δ4^*-and *blm^Δ11/Δ11^*-derived cells (**Figure 3f**). Our data suggests that canonical cellular mechanisms, including micronuclei and ATM-dependent H2AX phosphorylation, are conserved in killifish. We next sought to develop experimental strategies to ameliorate disease phenotypes.

### Genetic perturbation of the killifish cGAS–STING pathway

While the engraftment assay highlighted a possible role for immunity in A-T cancer predisposition, we were curious to explore the extent to which immune functions contribute to other disease phenotypes *in vivo.* To this end, we decided to specifically alter innate immunity by disrupting the cGAS-STING pathway. Using the strategies described above we mutated the *cgas* gene, producing homozygous mutants for the *cgas^Δ10/Δ10^*deletions (**Figure 4a**).

**Figure 4:**
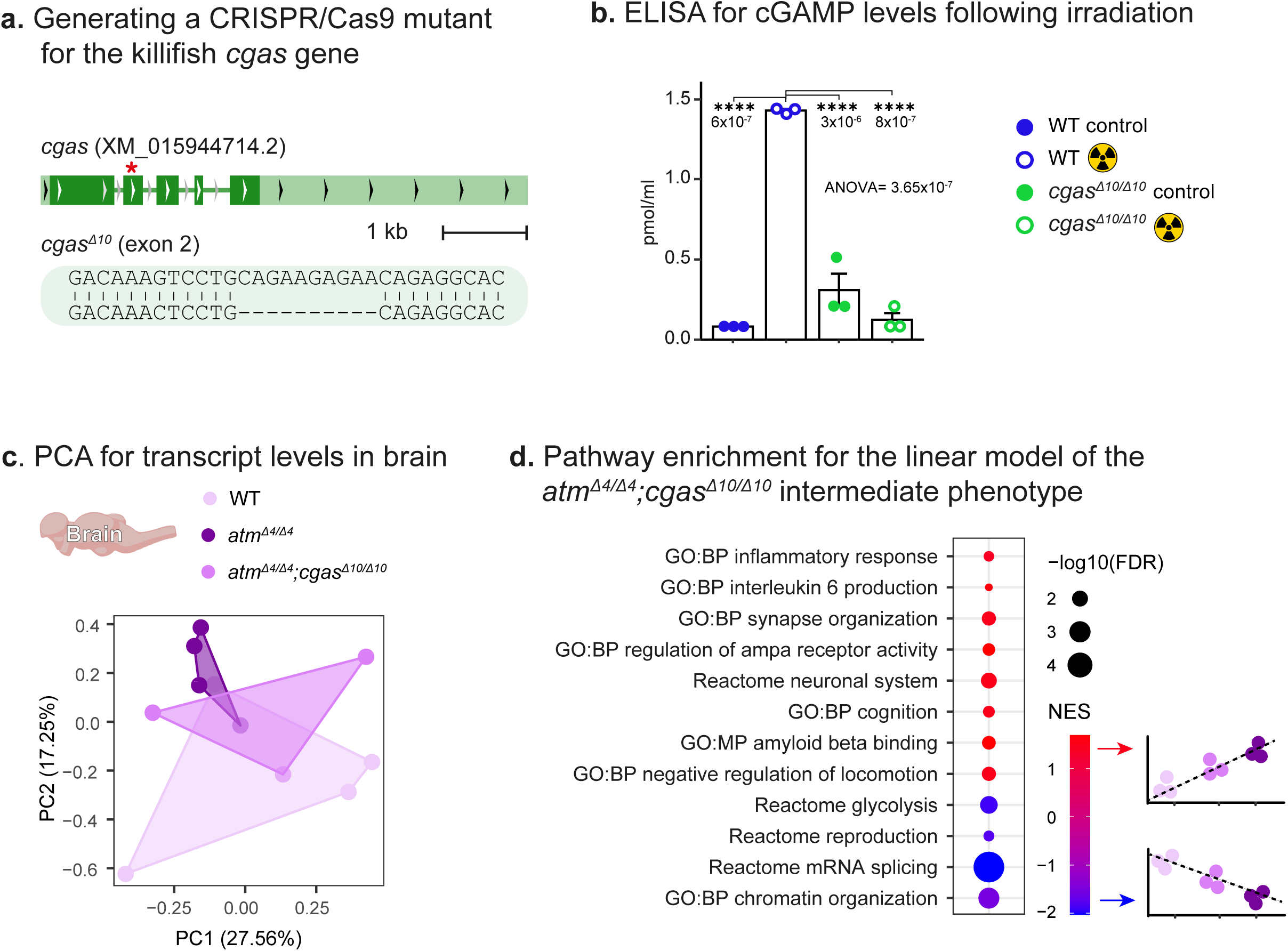
Transcriptional characterization of A-T brains following *cgas* inactivation. **(a)** Generation of a CRISPR knockout mutant for the *cgas* gene, showing the genomic locus with the mutation site (red asterisk) and a 10 base-pair deletion. **(b)** Following an irradiation treatment, cGAMP levels were calculated using an ELISA, in either WT or cgas^Δ10/Δ10^-derived primary fibroblast cultures. Significance was calculated using one-way ANOVA with Tukey post hoc, and *p* values are indicated. Error bars show mean ± SEM. **(c)** PCA of brain transcript levels, including WT, *atm ^Δ4/Δ4^,* and *atm ^Δ4/Δ4^;cgas ^Δ10/Δ10^* fish. *n* = 3-4 samples per condition. Each symbol represents an individual fish. **(d)** Dot plot showing functional enrichments (GO, FDR < 5%) using GSEA for differential gene expression of a linear model between WT, *atm^Δ4/Δ4^,* and *atm ^Δ4/Δ4^;cgas ^Δ10/Δ10^* fish. NES: normalized enrichment score.

To assess whether the *Δ10* indel is a loss-of-function allele, we quantified the levels of 2’3’-cyclic GAMP (cGAMP). The cGAMP cyclic dinucleotide second messenger is produced by cGAS following its binding to double-strand DNA (dsDNA). cGAMP then triggers STING activation, which initiates a robust inflammatory response through the type I interferon (IFN) pathway^15^. Quantification of cGAMP using ELISA revealed that *cgas^Δ10/Δ10^*-derived cells failed to synthesize cGAMP following ionizing irradiation-mediated DNA damage (5Gy, **Figure 4b**). Thus, our data indicates that the *cgas ^Δ10^* allele causes a loss-of-function.

### Transcriptional analysis suggests that cGAS perturbation might promote healthspan in A-T

As disease phenotypes are more severe in A-T fish, we selected this model to explore the role of the cGAS-STING pathway as a modifier of DDR syndromes. We focused on healthspan parameters, as the effect on lifespan was relatively mild in A-T (**Figures 2**, **S2**).

We first generated *cgas^Δ10/Δ10^*and *atm^Δ4/Δ4^* double mutants, and performed RNA sequencing in the brains and livers of the three genotypes: WT, *atm^Δ4/Δ4^*, and *atm^Δ4/Δ4^;cgas^Δ10/Δ10^*. Using PCA analysis, we observed that the brain samples segregate according to PC2, with the double mutants positioned between the WT and *atm^Δ4/Δ4^* samples (**Figures 4c**, **S4a**). We, therefore, performed a linear model to test which pathways are associated with the predicted intermediate phenotype of the double mutant brains (using sex as a covariant in the linear model, see **Methods**).

Using a gene set enrichment analysis (GSEA^60^, **Figures 4d**, **S4b**), enrichments that fit the linear model in the brain were linked with the diverse spectrum of A-T disease phenotypes^10^ (**Figure 4d**). Specifically, these pathways suggested that classical A-T pathologies, including inflammatory responses, epigenetic alterations, as well as modified brain, metabolic, and reproductive functions, could be partially ameliorated following cGAS inactivation. Therefore, to assess the direct effect on A-T pathophysiology, we explored selected disease phenotypes in the double mutants.

### cGAS perturbation ameliorates A-T pathology

Both male and female *atm^Δ4/Δ4^* fish are largely infertile and suffer from a severe germline defect (**Figure 2a, b**). To characterize germline developmental stages at higher resolution, we performed a single-molecule fluorescent in-situ hybridization chain reaction (smFISH HCR) in gonadal sections from mature (5-week-old) fish. We used stage-and sex-specific germline markers that we have recently identified^41^, including the spermatocyte marker *dmc1* (DNA meiotic recombinase 1) and spermatid marker *tekt1* (tektin-1) in males, and the pan germline marker *ddx4/vasa* (dead-box helicase 4) in females (**Figures 5a**, **S5a**). As a general marker of gonadal support cells in both sexes, we used *amh*^41^ (anti-mullerian hormone).

**Figure 5:**
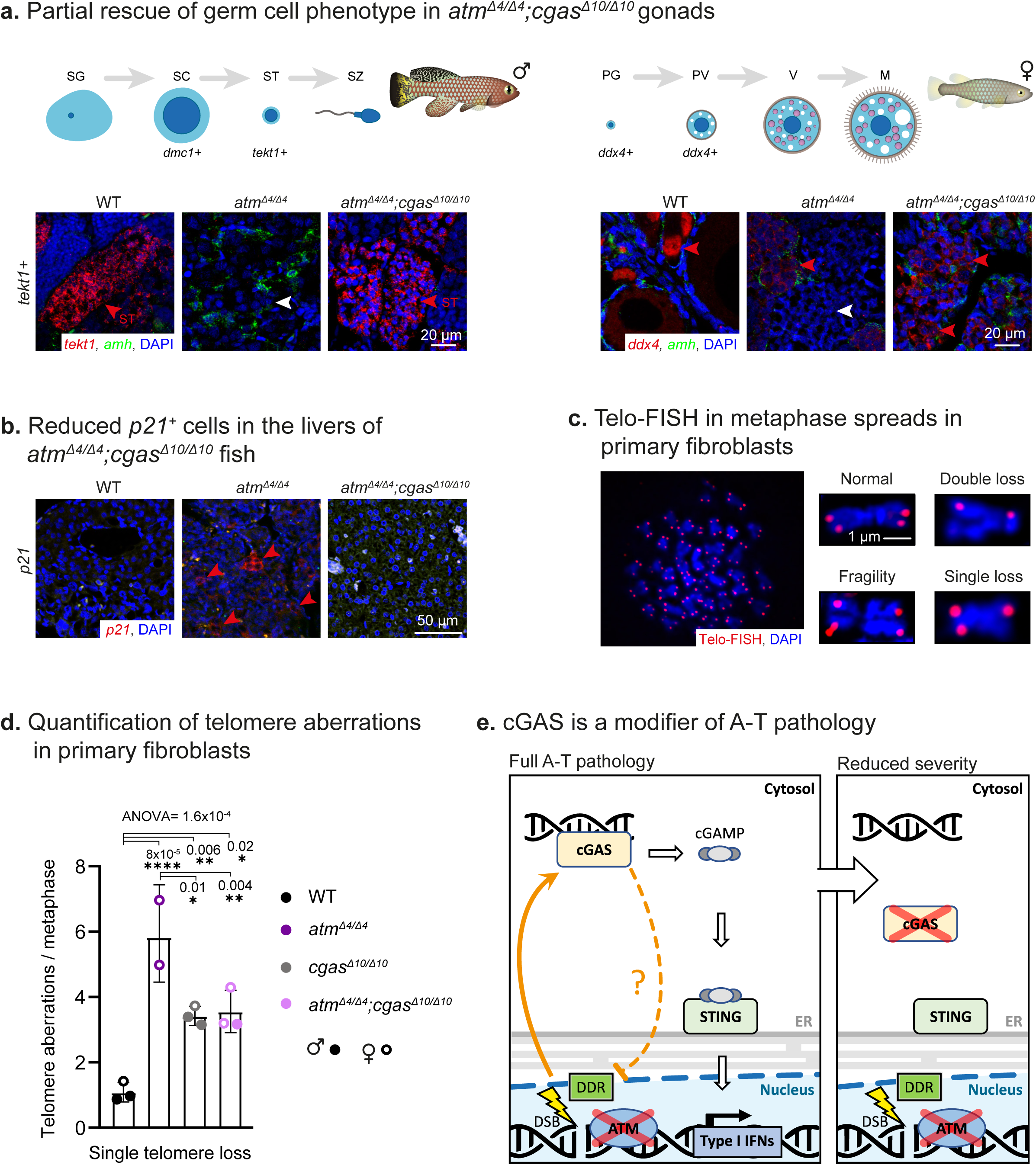
A-T-related physiological and cellular phenotypes following *cgas* inactivation. **(a)** Top: germ cell development in male and female killifish, adapted from^41^. Bottom: smFISH for germ cell markers in the testis (*tekt1*) and ovary (*ddx4*), in either WT, *atm^Δ4/Δ4^*, or *atm^Δ4/Δ4^;cgas ^Δ10/Δ10^* fish (red). The supporting cell marker *amh* is in green. Scale bar: 20 µm **(b)** smFISH for the senescence marker *p21* (red) in the liver of either WT, *atm^Δ4/Δ4^*, or *atm^Δ4/Δ4^;cgas ^Δ10/Δ10^* fish. Representative of *n* ≥ 3 mature (5-week-old) individuals. Scale bar: 50 µm **(c)** Detection of telomere aberration by fluorescence in situ hybridization (Telo-FISH, in red) in metaphase spreads from primary killifish fibroblast. **(d)** quantification of single telomere loss in WT, *atm^Δ4/Δ4^*,*cgas^Δ10/Δ10^*, and *atm^Δ4/Δ4^;cgas ^Δ10/Δ10^* primary killifish fibroblasts (derived from the indicated sexes). Significance was calculated using one-way ANOVA with Tukey post hoc, and *p* values are indicated. Error bars show mean ± SEM. **(e)** A proposed cellular mechanism for the amelioration of A-T disease phenotypes via cGAS inactivation For all smFISH experiments, we present representative images from at least 4 sections, derived from 3-6 mature (5-week-old) individuals

The smFISH staining suggests that the differentiated *tekt1^+^*spermatids were completely missing in *atm^Δ4/Δ4^* mutant males, while undifferentiated *dmc1^+^* spermatocytes were present in the testis of both WT and *atm^Δ4/Δ4^*males (*tekt1* in **Figures 5a***, dmc1* in **S5a,** n=3/3). Similarly, only immature oocytes at primary growth (PG) were visible in *atm^Δ4/Δ4^* ovaries (n=3/3, **Figures 5a**). Remarkably, performing these experiments in *atm^Δ4/Δ4^*;*cgas^Δ10/Δ10^* double-mutants revealed the presence of *tekt1^+^*spermatids (n=5/5), and an expansion of *ddx4/vasa* expression in the ovaries (n=6/6, **Figure 5a**). H&E staining further identified mature spermatozoa in the double-mutant testis of young fish (5-week-old), which persisted to old age (15-week-old, **Figure S5a**).

To explore the impact on cellular senescence, another progeroid A-T pehnotype^10^, we performed smFISH in liver sections for the senescence marker^61^ *cyclin-dependent kinase inhibitor 1a* (*cdkn1a*, or *p21*). Our findings suggest that while *p21^+^* cells are detected in *atm^Δ4/Δ4^* fish, they are no longer visible in *atm^Δ4/Δ4^*;*cgas^Δ10/Δ10^* double-mutants (n≥4 for each group, **Figure 5b**).

### cGAS perturbation positively affects genomic stability in A-T

So far, we have designed our experiments based on the assumption that cGAS activation occurs downstream of genomic instability. However, we were curious to determine whether classical markers of genomic instability, including telomere aberrations and de-repression of transposable element (TEs) were affected in the double mutants.

To characterize the nature of telomere aberrations, we examined individual telomeres by FISH (Telo-FISH^62^). This sensitive approach can detect various aberrations, including telomere fragility, single-loss, and double-loss of telomeres. To this end, we performed Telo-FISH on metaphase chromosomes in primary cells derived from the different genotypes (**Figure 5c**, **d**, see **Methods** for the modified killifish protocol). Our data suggest that while *atm^Δ4/Δ4^*-derived cells display a robust increase in aberrations, *atm^Δ4/Δ4^;cgas^Δ10/Δ10^* double mutants were partially rescued (**Figures 5d**, **S5b**). Notably, the loss of cGAS alone can also contribute to a mild increase in aberrations. Consistent with our previous findings^42^, ex vivo primary cells derived from either males or females display comparable phenotypes once cultured (**Figures 5d**, **S5b**).

To further investigate the global effect on TE expression, we reanalyzed our PolyA-selected transcriptomic data. While many transposons utilize host Pol II machinery for transcription^63^, others can still be assessed due to their abundance and multi-copy nature. Our analysis revealed that A-T fish exhibited global reactivation of TE transcripts, particularly in female brains (**Figure S5c**), while double-mutants showed a rescue to WT levels. A multi-organ transcriptomic dataset from aging male and female killifish^64^ provides some insights into these sexually dimorphic responses. Notably, cytokine and Type I interferon production are innately higher in females, especially in the brain (**Figure S5d**).

In conclusion, our findings suggest that blocking the cGAS-STING axis can rescue specific A-T disease phenotypes *in vivo*, highlighting a key role of immune functions in disease progression. These data may have broader implications for A-T treatment through pharmacological cGAS inhibition, which will require further investigation to minimize possible adverse effects (e.g. telomere aberrations). Furthermore, our findings suggest sexually dimorphic mechanisms of DDR, and can serve as a paradigm for other DDR syndromes (see **Figure 5e**).

## Discussion

As individuals age, the immune system undergoes many functional alterations^4,16^, including low-level chronic inflammation (‘inflammaging’^19^), shifts in immune cell composition (e.g., the ’myeloid shift’), and reduced adaptive responses. These changes are influenced by a variety of factors, including DNA damage accumulation^19^ (as discussed above), chronic viral infections (e.g., CMV^65^), protein aggregate buildup^66^, secretion of pro-inflammatory cytokine by senescent cells^67^, and hematopoietic stem cell depletion^68^. Ultimately, this decline can result in heightened susceptibility to infections and diminished vaccination responses.

Inflammaging can also exacerbate other age-related pathologies^16,69,70^, such as interfering with the clearance of precancerous cells^71^. However, the precise mechanisms underlying inflammaging, its overall impact on aging and DDR-related diseases, and the potential for reversing these pathologies remain to be elucidated. To explore these questions, we generate genetic killifish models for the BS and A-T, which replicate classical disease phenotypes. We then investigate the contribution of the cGAS-STING pathway to A-T disease pathophysiology, and demonstrate that *atm^Δ4/Δ4^;cgas^Δ10/Δ10^* double mutants exhibit attenuated disease progression. Thus, proposing the cGAS–STING pathway as an important driver of A-T.

### DNA repair across evolutionary distances

In support of these findings is the importance of DNA repair mechanisms in comparative evolution, which suggests that the DDR has coevolved with vertebrate longevity^72–80^. For example, mammalian genomic studies revealed that the long-lived naked mole-rat displays enhanced resistance to DNA damage^81^, and an increase in *tp53* copy number is linked with cancer resistance in elephants^78^. Comparative transcriptomic studies also demonstrated similar trends in many long-lived mammalian species^72,73^.

Similarly, in fish, a genomic study of 45 killifish species has suggested that DNA repair genes are under relaxed selection in short-lived species^75^, while positive selection for DNA repair pathways was observed in the long-lived rockfish^76,80^ and the Greenland shark^82^. Finally, the longevity of bats has been proposed to be associated with a diminished ability to sense DNA damage^83,84^. While the mechanism proposed in bats might seem counterintuitive, it could potentially lead to a protective decrease in age-related sterile inflammation^19^ (’inflammaging’), naturally mimicking an optimized inhibition of the cGAS-STING pathway.

### cGAS may suppress the DDR

The cGAS protein is traditionally known as the primary receptor for cytosolic DNA, promoting type I interferon responses^85^. However, recent studies have shown that cGAS can also be tethered to chromatin in the absence of external stimuli^86–88^. Interestingly, evidence suggests that in the nucleus, cGAS may inhibit DNA repair, particularly through homologous recombination (HR)^89^. Our findings support this seemingly counterintuitive phenomenon, suggesting that mutating cGAS in the context of A-T can improve telomere aberrations.

One proposed mechanism is that DNA damage triggers interaction with activated Poly ADP-ribose polymerase 1 (PARP1), which prevents the recruitment of proteins necessary for HR^90^. Another possibility is that chromatin-bound cGAS inhibits the formation of displacement loops, a critical step for HR^91^. As a result, cells expressing cGAS show increased accumulation of double-strand breaks (DSBs) following irradiation compared to cGAS-deficient cells.

Notably, this non-canonical cGAS function appears independent of cGAS-mediated innate immune sensing^91^. Conversely, the increase in telomere damage following the loss of cGAS alone could be mediated via an unknown mechanism, or through the inhibition of cell death^92^ (allowing damaged cells to persist). Therefore, specifically mutating the enzymatic activity of cGAS may be required for future studies to uncouple these mechanisms, thus allowing for the design of tailored interventions that minimize adverse outcomes.

---

**Figure S1:**
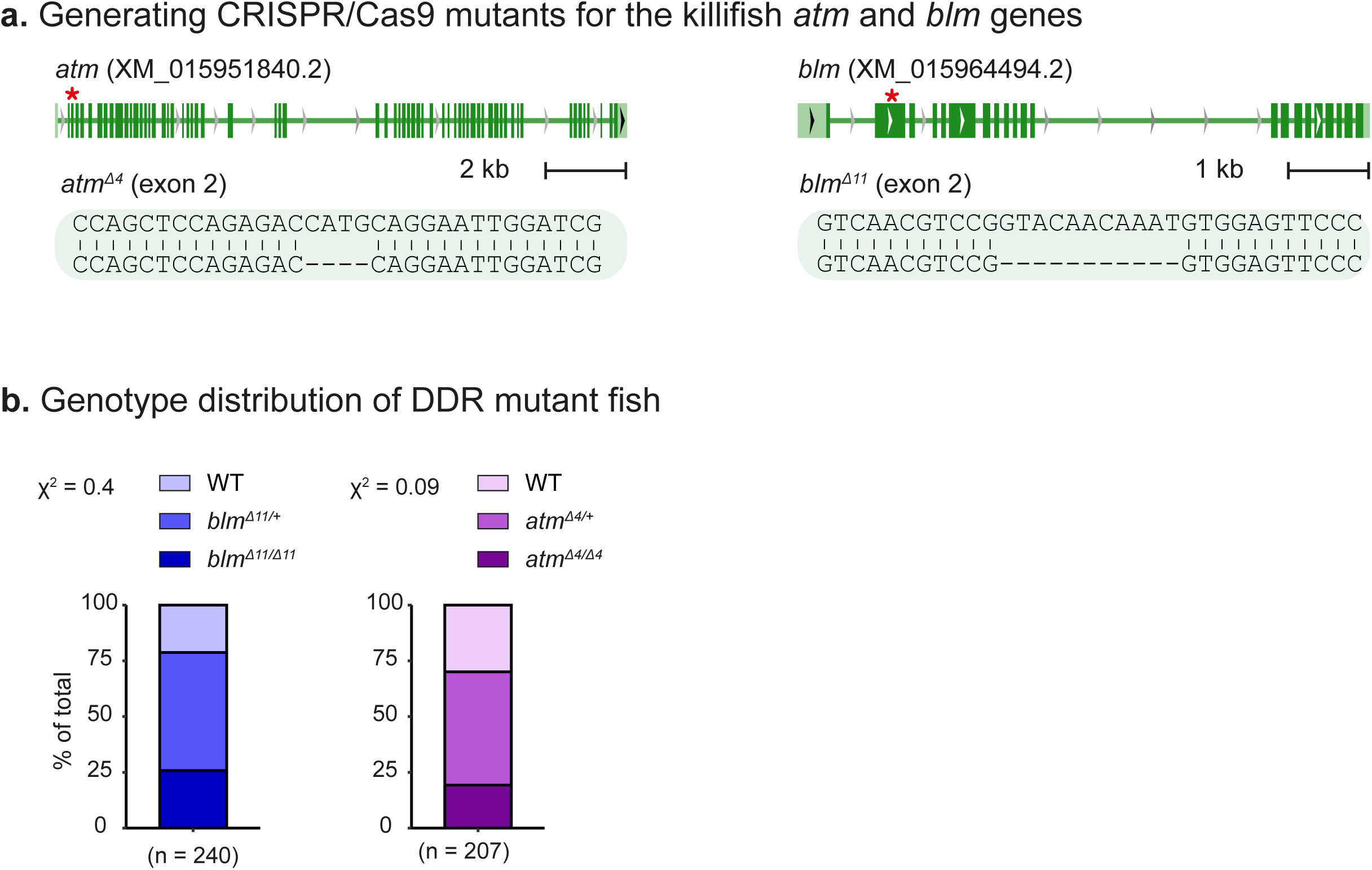
Generating killifish models for Ataxia-Telangiectasia and Bloom Syndrome. **(a)** Generation of CRISPR mutants for the *atm* and *blm* genes, showing the gRNA target (red asterisk), and resulting delusions. **(b)** Distribution of genotype progeny from heterozygous pairs. *n* = 207-240 individuals per genotype. Significance was measured by a two-sided χ^2^ test with Mendelian proportions (25:50:25) as the expected model.

**Figure S2:**
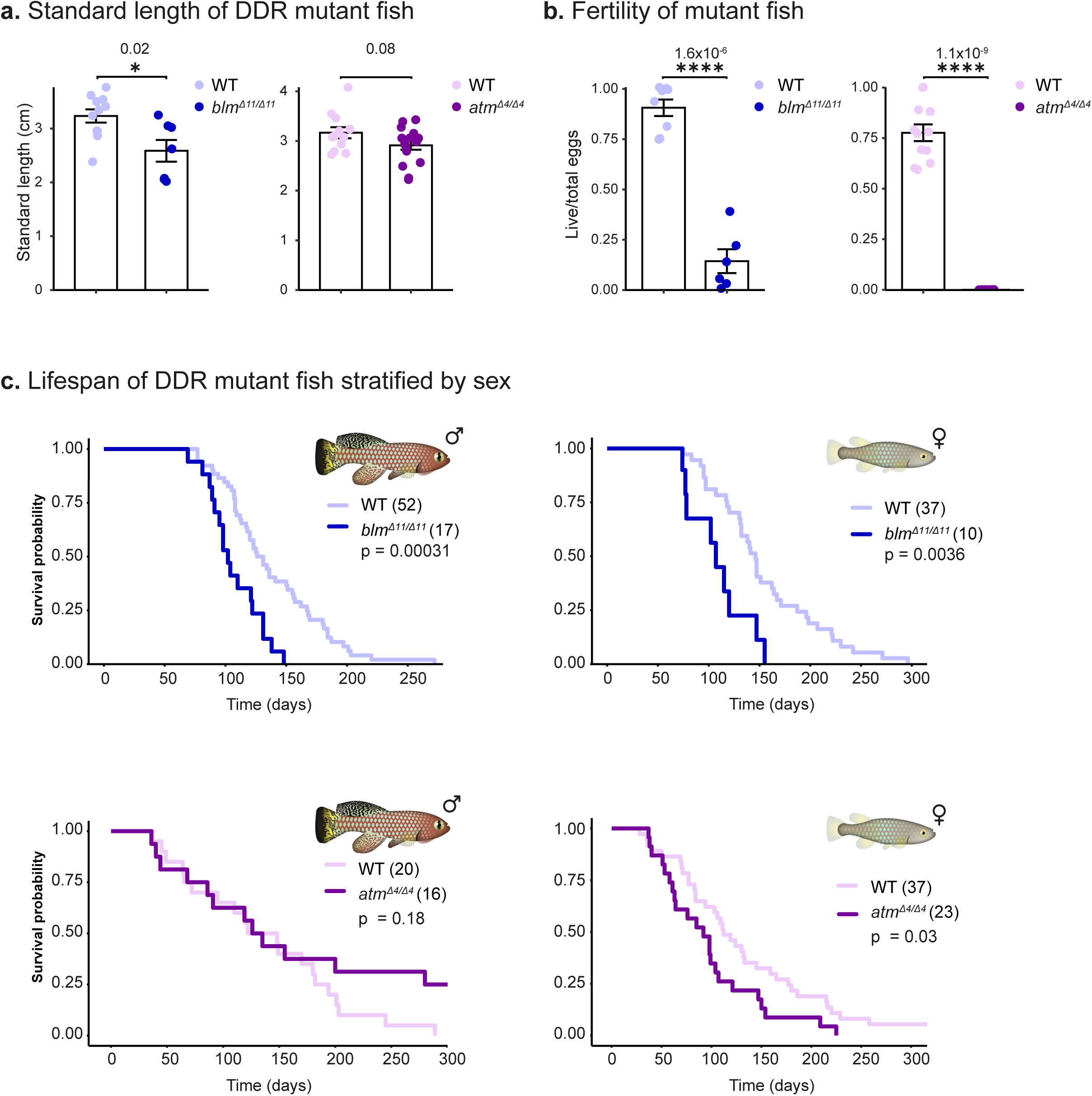
Assessing fish size and fertility in the killifish DDR mutants. **(a)** Standard length measurements of ∼15-week-old male fish. *n* ≥ 7 fish per group. Significance was calculated using an unpaired t-test. Error bars show mean ± SEM. Exact p-values are indicated. **(b)** Quantification of female fertility output. Each dot represents the ratio between live and total eggs for the indicated genotypes. Error bars show mean ± SEM. Significance was calculated using an unpaired t-test, and *p* values are displayed. **(c)** Lifespan of WT, *atm^Δ4/Δ4^*, and *blm^Δ11/Δ11^* fish, assessed separately for male and female. *p* values for differential survival in log-rank tests and fish numbers are indicated.

**Figure S3:**
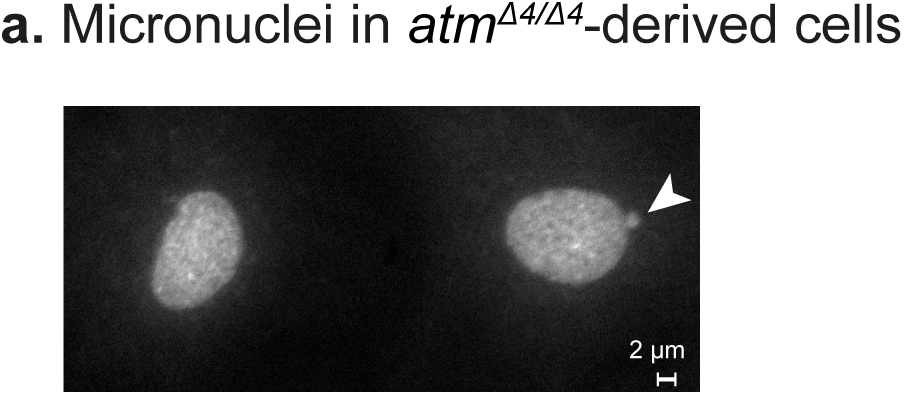
**(a)** Micrographs of DAPI stained *atm^Δ4/Δ4^*-derived primary fibroblasts. A cell with a micronucleus is shown with a white arrow.

**Figure S4:**
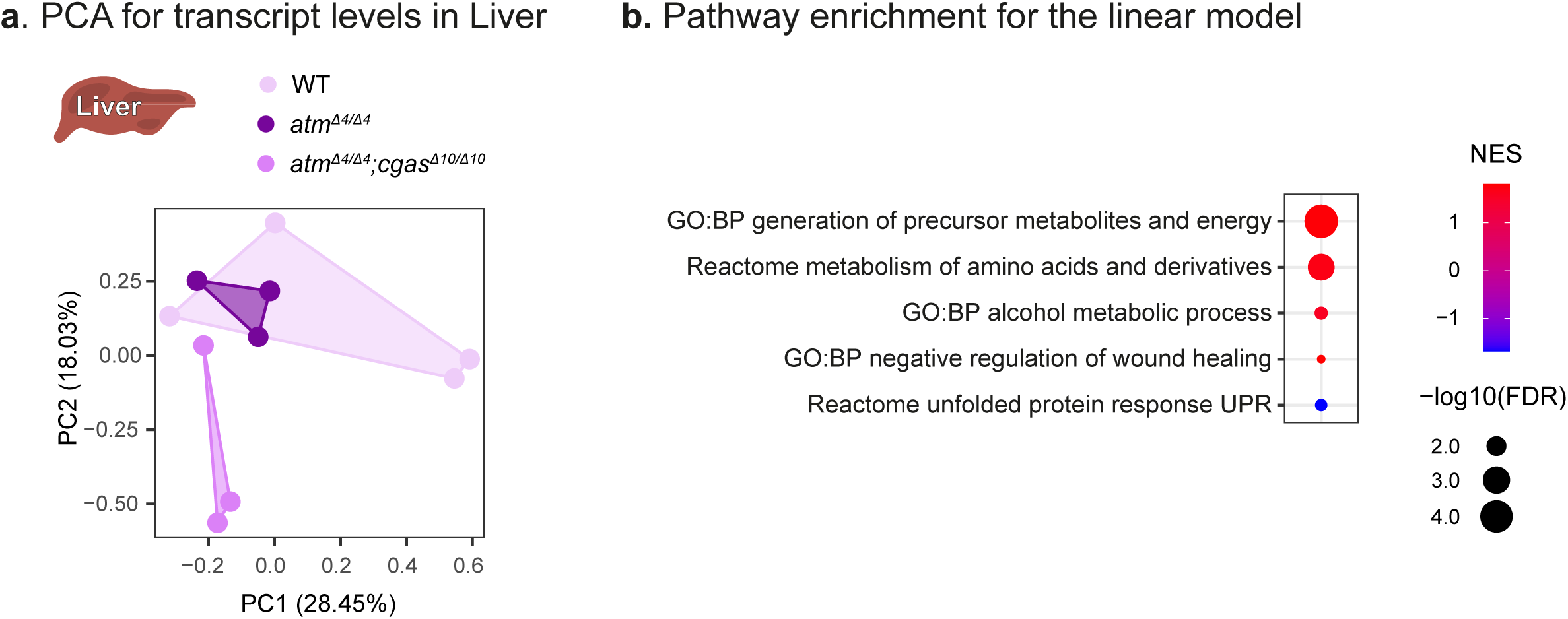
Transcriptional characterization of A-T livers following *cgas* inactivation. **(a)** PCA of liver transcript levels, including WT, *atm ^Δ4/Δ4^,* and *atm ^Δ4/Δ4^;cgas ^Δ10/Δ10^* fish. *n* = 3-4 samples per condition. Each symbol represents an individual fish. **(b)** Dot plot showing functional enrichments (GO, FDR < 5%) using GSEA for differential gene expression in a linear model (see Figure 4d). NES: normalized enrichment score.

**Figure S5:**
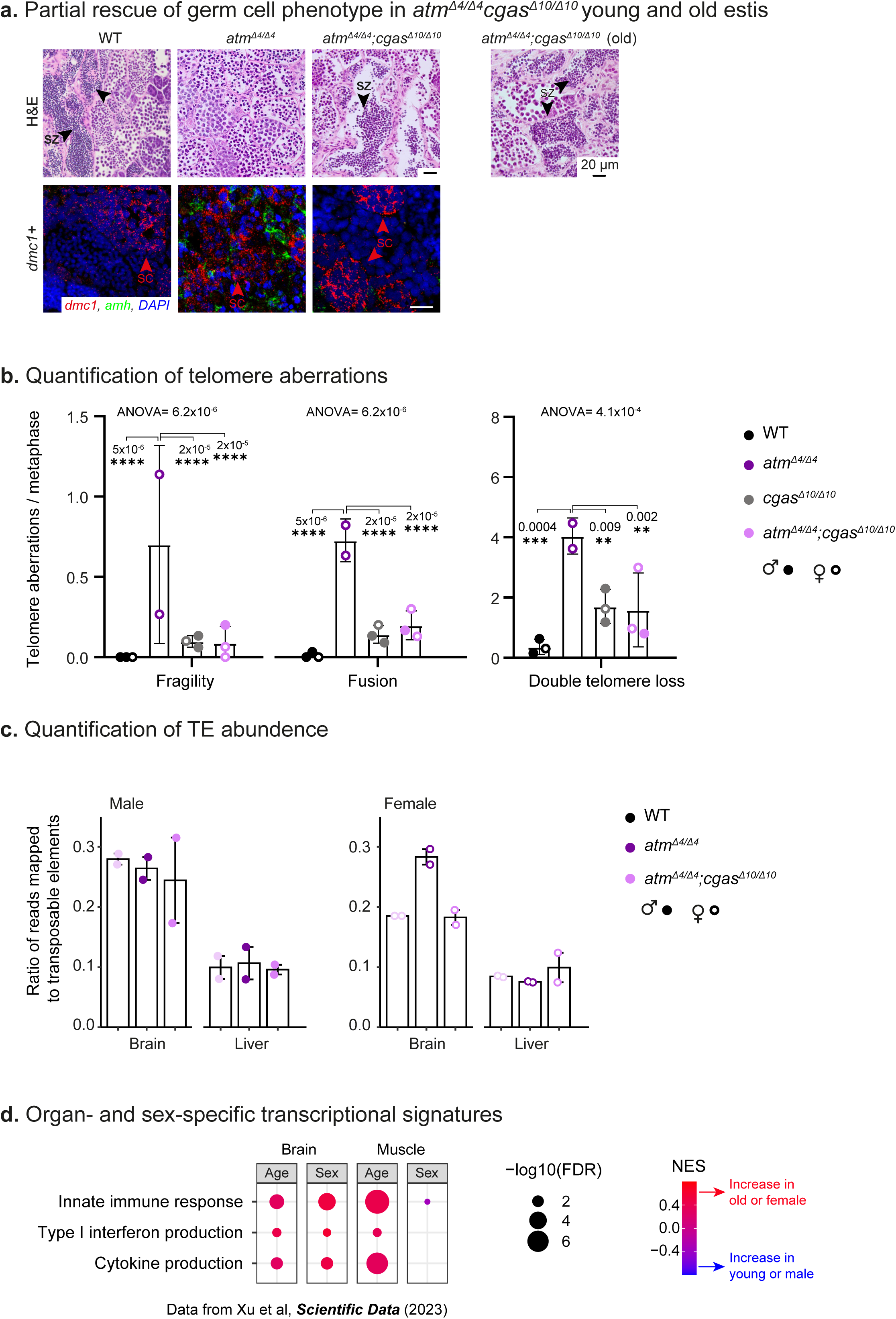
Germline pathologies in Ataxia-Telangiectasia are improved following *cgas* inactivation. **(a)** Left: H&E (top) and smFISH (bottom) for immature germ cell marker in the testis (*dmc1*), in young (5-week-old) WT, *atm^Δ4/Δ4^*,and *atm^Δ4/Δ4^;cgas ^Δ10/Δ10^* fish (red). The supporting cell marker *amh* is in green. Right: H&E in old (15-week-old) *atm^Δ4/Δ4^;cgas ^Δ10/Δ10^*male. Representative of *n* ≥ 3 individuals. Scale bar: 20 µm **(b)** Quantification of telomere fragility, telomere fusion, and double telomere loss in WT, *atm^Δ4/Δ4^*,*cgas^Δ10/Δ10^*, and *atm^Δ4/Δ4^;cgas ^Δ10/Δ10^* primary killifish fibroblasts (derived from the indicated sexes). Significance was calculated using one-way ANOVA with Tukey post hoc, and *P* values are indicated. Error bars show mean ± SEM. **(c)** Quantification of the ratio of transcripts mapped to TEs in the liver and brain data (Figures 4, S4), of WT, *atm^Δ4/Δ4^*, and *atm^Δ4/Δ4^;cgas ^Δ10/Δ10^*fish stratified by sex. **(d)** Dot plot showing functional enrichments (GO, FDR < 5%) using GSEA for differential gene expression for a multi-organ transcriptomic killifish data^64^ (comparing either between ages or sexes). Enrichments reveal sexual dimorphism of key pathways linked to innate immunity and interferon type I. NES: normalized enrichment score.

## Acknowledgments

We thank E. Cohen, M. Goldberg and the Harel laboratory for stimulating discussion and feedback on the manuscript. We thank A. Abu-tair, Y. Birenbaum, F. Idrees and R. Barakat for help with killifish maintenance and N. Melamed-Book from the imaging core (HUJI). This study was supported by ERC StG no. 101078188 (I.H.), the Zuckerman Program (I.H.), the ISF 2178/19 (I.H.), Israeli Ministry of Science 3-17631 (I.H.) and 3-16872 (I.H.) grants, the Moore Foundation GBMF9341 (I.H.), BSF-NSF 2020611 (I.H.), NIH 1R21AG063739 (I.H. and B.A.B.), KAMLA Research Fund of the Hebrew University of Jerusalem and Sha’are Zedek Medical Centre (M.B.), the Israeli Ministry of Agriculture 12-16-0010 (I.H.), the Levi Eshkol Scholarship of the Israeli Ministry of Science (E.M.), the Pamela and Paul Austin Research Centre on Aging fellowship (T.A.). R35GM142395 and R35GM142395-03S1(B.A.B.).

## Contributions

M.B. and U.G. designed the study and performed the experiments, under the supervision of I.H. and I.B.. G.A. generated the *atm* mutants, U.G. generated the *cgas* and *blm* mutants. M.B. performed the histology and smFISH, with help from E.M. T.A. designed and performed the computational analysis for the RNA sequencing, under the supervision of I.H.. R.H. and H.E. performed the metaphase spread preparation and Telo-FISH under the supervision of Y.T. A.J.J.L. analyzed TE expression, under the supervision of B.A.B.. T.A. and U.G. performed statistical analyses. M.B., U.G., T.A and I.H. wrote the manuscript. All authors commented on the manuscript.

## Ethics declarations

The authors declare no competing interests.

## Methods

### EXPERIMENTAL MODELS

#### African turquoise killifish strain, husbandry, and maintenance

The African turquoise killifish (GRZ strain) was housed as previously described^41,42^ at 28°C in a central filtration recirculating system, with a 12 h light/dark cycle at the Hebrew University of Jerusalem (Aquazone ltd, Israel). Until the age of 2 weeks, fish were exclusively fed with live Artemia (#109448, Primo). Starting week 3, fish were fed three times a day on weekdays (and once a day on weekends), with GEMMA Micro 300 Fish Diet (Skretting Zebrafish, USA), supplemented with Artemia twice a day. In these conditions, killifish lifespan was approximately 4-6 months. Mutant alleles were maintained as heterozygous and propagated by crossing with wild-type fish. For experiments heterozygotes were paired to generate homozygotes, the resulting offspring were not used for propagation. All turquoise killifish care and uses were approved by the Subcommittee on Research Animal Care at the Hebrew University of Jerusalem (IACUC protocols #NS-18-15397-2, #HU-24-17607-4).

#### CRISPR/Cas9 target prediction and gRNA synthesis

CRISPR/Cas9 genome-editing protocols were performed as previously described^41,42^. In brief, we targeted conserved regions that are upstream to functional or active protein domains. Target sites were identified using Synthego (https://www.synthego.com/products/bioinformatics/crispr-design-tool) for *blm*, or CHOPCHOP (https://chopchop.rc.fas.harvard.edu/)^93^ for *atm* and *cgas.* gRNA sequences were as follows (PAM sites are in bold):

*blm* exon 2: 5’: GGACTCCACATTTGTTGTAC**CGG**-3’;

*atm* exon 3: 5’: AGGTGCGATCCAATTCCTGCA**TGG**-3’;

*cgas* exon 2: 5’: GGCATTGAAACGTGATCCAAC**TGG**-3’.

Design and hybridization of variable oligonucleotides (which are gRNA-specific) with a universal reverse oligonucleotide was performed according to^41,42^, and the resulting products were used as a template for in vitro transcription. gRNAs were in vitro transcribed and purified using the TranscripAid T7 kit (ThermoFisher, # K0441), according to the manufacturer’s protocol.

#### Production of Cas9 mRNA

Experiments were performed according to^41,42^. The pCS2-nCas9n expression vector was used to produce Cas9 mRNA (Addgene, #47929). Capped and polyadenylated Cas9 mRNA was *in-vitro* transcribed and purified using the mMESSAGE mMACHINE SP6 ULTRA (ThermoFisher # AM1340).

#### Microinjection of turquoise killifish embryos

Microinjection of turquoise killifish embryos was performed according to^41,42^. Briefly, nCas9n-encoding mRNA (300 ng/μL) and gRNA (30 ng/μL) were mixed with phenol red (P0290, Sigma-Aldrich) and co-injected into one-or two-cell stage fish embryos. Sanger DNA sequencing was used for detecting successful germline transmission on F1 embryos. Fish with desired alleles were maintained as stable lines and further outcrossed to minimize potential off-target effects. The genomic area encompassing the targeted site (∼600 bp) was PCR-amplified using the following primer sequences:

BLM_F: 5’-aaaacacaaacactttgccctgt-3’; BLM_R: 5’-cagactttgctaaaggagtctga-3’;

ATM_F: 5’-ctgagggtgtggtctgatagc -3’; ATM_R: 5’-tctccttctggaggtaacgc-3’;

cGAS_F: 5’-ttaaggaaccccttcgcact-3’; cGAS_R: 5’-tggtaagcataaacatgctgcct-3’;

### METHOD DETAILS

#### Organ isolation

Individual killifish, according to the specified genotype, were sedated in 200 mg/L and then euthanized in 500 mg/L of Tricaine in system water. Animals were dissected under a binocular stereo microscope (Leica S9E) according to^45^. Whole brains and whole livers were harvested, and placed in separate tubes. Tubes were immediately snap-frozen in liquid nitrogen, and stored in −80°C until all samples were collected. All samples were collected during the morning time, between 7 am to 10 am, before morning feeding, to reduce the potential confounding effects driven by circadian rhythms. Age and sex of each genotype is included in **Table S1.**

#### RNA sequencing

##### RNA-seq library preparation

Organs were isolated as described above. Samples were disrupted by bead beating in 600μl of TriReagent (Sigma, T9424) and a single 3 mm metal bead (Eldan, BL6693003000) using TissueLyzer LT (QIAGEN, #85600) with a dedicated adaptor (QIAGEN, #69980). Beating was performed twice at 50 Hz for 2 min. RNA extraction was performed with Direct-zol RNA Purification Kits (Zymo). RNA concentration and quality were determined by using a Thermofisher Nanodrop One (ThermoFisher, ND-ONE-W) and Agilent 2100 bioanalyser (Agilent Technologies), respectively. Library preparation was performed using KAPA mRNA HyperPrep Kit (ROCHE-08105936001, according to the recommended protocols). Library concentrations were measured by Qubit (dsDNA HS, Q32854), and quality was measured by Tape Station (HS, 5067-5584). Libraries were sequenced by NextSeq 2000 P3, 50 cycles, 70 bp single-end (Illumina, 20046810) with ∼35 million reads per sample.

##### RNA sequencing analysis

Quality control and adapter trimming of the fastq sequence files were performed with FastQC (v0.11.8)^94^, multiQC (v1.12)^95^, fastx-toolkits (v0.0.13), Trim Galore! (v0.6.4)^96^, and Cutadapt (v3.4) ^97^. Options were set to remove Illumina TruSeq adapters and end sequences to retain high-quality bases with *phred* score > 20 and a remaining length > 20 bp. Successful processing was verified by re-running FastQC. Reads were mapped and quantified to the killifish genome Nfu_20140520^38,98^ using STAR 2.7.6a^99^. One fish from the *blm* analysis (B9) and one fish from *atm-cgas* analysis (Male-KO-4) were removed due to inflammatory phenotypes such as an enlarged spleen. Differential gene expression was performed using the edgeR package (v3.32.1)^100,101^. The analysis between *blm* mutant and WT fish using the classic model (exactTest function). The analysis of the WT, *atm* mutant, and *atm;cgas* mutants was performed using a linear model (glmQLFTest function), placing the *atm;cgas* mutant in between WT and *atm,* including a sex as co-variance. The ComBat_seq function from the sva package (v3.42.0)^102^ was used for batch correction.

##### Gene Ontology Enrichment Analysis

Enriched Gene Ontology (GO) terms associated with transcript levels (from the analysis mentioned above) were identified using either Gene Set Enrichment Analysis (GSEA) or GO enrichment analysis implemented in R package clusterProfiler (v3.18.1)^103^. In GSEA, all the transcripts were ranked and sorted in descending order based on the multiplication of log2 transformed fold change and -log10(FDR). Note that due to the random seeding effect in GSEA, the exact p-value and rank of the enriched terms may differ for each run. These random seeds did not qualitatively affect the enrichment analyses. In GO enrichment analysis, the threshold of FDR is >0.01, and the log2(FC) threshold is <1. GO terms were based on human GO annotations from org.Hs.eg.db (v3.13.0)^104^ and AnnotationDbi (v1.54.1)^105^.

##### Principal component analysis (PCA)

Standardized log2 transformed normalized count per million (CPM) was used as input for PCA. PCA was performed using autoplot function implemented in R package ggfortify (v0.4.12) and plotted using ggplot2 (v3.3.5).

#### Transposable element read ratio

Raw single-end FASTQ RNA-seq files were trimmed of adapters and low-quality reads were filtered using fastp version 0.23.4, using parameter “--adapter_sequence= AGATCGGAAGAGCACACGTCTGAACTCCAGTCAC”. Trimmed QC reads reads were mapped to the killifish reference genome (GCA_014300015.1) using STAR version 2.7.11b^99^ with the following parameters: --outFilterMultimapNmax 200, --outFilterIntronMotifs RemoveNoncanonicalUnannotated, --alignEndsProtrude 10 ConcordantPair, --limitGenomeGenerateRAM 60000000000, and --outSAMtype BAM SortedByCoordinate. Gene and TE count matrices were generated from aligned BAM files together with the killifish reference gene annotation and TE annotation as previously described^106^ using TEtranscripts version 2.2.1^107^. TEtranscripts count matrices were then used to estimate both overall gene and TE expression levels by integrating existing gene annotations (GTF) with a custom TE annotation (GTF) based on FishTEDB (see^64,106^). To determine the ratio of reads contributed by TE regions, count matrices generated by TEtranscripts were imported into R version 2023.03.0+386. The sum of reads mapped to TE features was then divided by the total sum of reads for each tissue sample, respectively to obtain the global change of TEs in the different treatments/sex/tissue.

#### Gene length analysis

Gene length (including the introns), was calculated manually using the GTF annotation file. In the gene length analysis, we calculated the correlation between gene length and the fold-change between WT and *blm* mutant, and displayed the distributions of the gene length in up-or down-regulated genes^17^.

#### Survival, Standard length, and fertility assays

##### Lifespan

For reproducible lifespan experiments, constant housing parameters are very important^45,108^. Following hatching, fish were raised with the following density control: 10 fish in a 1-liter tank for week 1, 5 fish in a 3-liter tank for weeks 2-4. From this point onwards, adult fish were genotyped and single-housed in 1-liter tanks for their remaining lifespan. Plastic plants were added for enrichment. Both male and female fish were used for lifespan experiments and were treated identically. Fish mortality was documented daily starting week 4. Lifespan analyses were performed with a Kaplan-Meier estimator and significance was calculated with a log-rank test with FDR adjustment, using the R programming language (4.1.3) and additional packages. The packages used were “survival” (v.1.3.1), and “survminer” (v.3.2.13) for analysis and graphing, and “readxl” (v:0.4.9) and “dplyr” (v:1.0.8) for reading input and arranging data.

##### Standard length

For measuring standard length, fish were imaged at the age of 13-15 weeks with a Canon Digital camera EOS 250D, prime lens Canon EF 40 mm f/2.8 STM that documented body length. To limit vertical movement during imaging the camera was mounted on a tripod, fish were positioned in a water tank with 3 cm water depth, and images were taken from the top using fixed lighting and height. A ruler was included in each image for an accurate scale. Body length was then calculated (using MATLAB R2021a), by converting pixel numbers to centimeters using the included reference ruler. Data was analyzed and plotted with R programming language (4.1.3). Significance was calculated using an unpaired t-test.

##### Fertility Analysis

Fish fertility was evaluated according to^41,42^. Briefly, Adult fish (6-7 weeks old) were paired as 1 male and 1 female. for *blm^Δ^*^11^*^/Δ11^*and *atm^Δ4/Δ4^* 3 homozygote and 3 WT pairs were placed each in a 3-liter tank. All breeding pairs were allowed to continuously breed on sand trays, and embryos were collected and counted weekly for 4 weeks. Unfertilized eggs are easily identified, as they die shortly after egg-laying and the yolk becomes opaque. Results were expressed as a ratio of fertilized eggs per week of egg-lay. Each egg collection from each pair was considered as one data point. Significance was calculated using an unpaired t-test with R programming language (4.1.3). Normality was tested with Shapiro-Wilk test and assessed with QQ-plots.

#### Histology

Tissue samples were processed as described previously^12,35–37,40,43,46,98,109–118^. Briefly, for paraffin sections, the body cavity of the fish was opened. Animals were fixed for 48h in 4% PFA solution in neutral pH PBS at 4°C. Samples were then washed and incubated in 0.8M EDTA with 1% PFA for 1 week for decalcification, then embedded in paraffin by standard procedures.

#### Fluorescence in-situ Hybridization

Fish were sacrificed and fixed as detailed above, and then embedded in paraffin by standard procedures. Paraffin sections (7-4 µm) were deparaffinized with Histo-Clear (Bar-Naor, 64110-01), rehydrated in a series of decreasing concentrations of ethanol solutions (15 minutes in 100% twice, 5 minutes in 90%, 70%, 30%), and finally water. Fluorescence *in-situ* hybridization was carried out as per manufacturer’s protocol, HCR™ RNA-FISH protocol (Molecular Instruments). The probes used were custom-designed by Molecular Instruments. Slices were imaged using the FV-1200 confocal microscope (Olympus, Japan). The objectives that were used: 40X/0.9. Multiple dyes sequential scanning mode was used to avoid emission bleed-through. Fluorophore (Excitation, Emission): DAPI (405nm, 430-470nm), Green (488 nm, 505-550 nm), Red (561 nm 570-620 nm). Images were then scored by hand using FIJI software pack for ImageJ for the area of the transcript of interest (i.e. *dmc1, tekt1, p21, ddx4*) out of the gonad area in the image. At least 4 sections from at least 3 male fish were scored for each staining.

#### Cell culture

##### primary fibroblast cultures from killifish tail fins

Adult fish (8-15 weeks old), of the indicated genotype and gender were sedated with MS-222 (200 mg/L Tricaine, in system water). All following experiments were conducted at 28°C unless stated otherwise. Following sedation, a 2–3 mm tissue was trimmed from the tail fin using a sterile razor blade and individually disinfected for 10 min with a 25-ppm iodine solution (PVP-I, Holland Moran 229471000) in Ringer solution (Sigma, 96724). Tissue samples were then incubated for about 2h with 1 mL of an antibiotic solution containing Gentamicin (50 µg/mL Gibco) and PrimocinTM (50 µg/mL, InvivoGen) in DPBS at room temperature. Tissues were then washed with sterile DPBS and transferred into an enzymatic digestion buffer (200 µL, in a 24-well plate) containing Dispase II (2 mg/mL, Sigma Aldrich) and Collagenase Type P (0.4 mg/mL, Merck Millipore) in Leibovitz’s L-15 Medium (Gibco). During the first 7 days, cells were washed daily with fresh media before adding new media. When cells reached 85– 90% confluency, they were passaged with Trypsin-EDTA 0.05% (0.25% Trypsin-EDTA, diluted in DPBS). Cells were incubated at 28°C humidified incubator (Binder, Thermo Scientific) with normal air composition, and were used for downstream applications between passages 6-10.

##### Proliferation assay

Cells from individual fish were seeded in a 35 mm dish, at a density of approximately 67,000 cells/plate. 24h and 120h after seeding, cells were counted using CellDrop automatic cell counter (DeNovix, CellDrop BF). The medium was not changed during the experiment. The experiment was performed twice using 3 biologically independent replicates for the cultures with the indicated genotypes (using the mean of four measurements per dish) in each experiment. Data was analyzed and plotted with R programming language (4.1.3). Significance was calculated using one-way ANOVA with Tukey post hoc.

##### Assessing γ-H2AX coverage using immunofluorescence

WT, *blm^Δ11/Δ11^* and *atm^Δ4/Δ4^* cells from at least 3 individual fish were treated with Etoposide, followed by immunostaining against γ-H2AX according to published protocols^41,42^. Briefly, cells from individual fish were seeded in 12 well plates at 40,000 cells/well with a coverslip and allowed to grow overnight. The next morning cells were treated with Etoposide (Sigma, E1383) at 50mM, or with DMSO at the same concentration for 1 hour. Media was removed, cells were washed once with sterile DPBS and fixed with 4% PFA for 15 minutes at room temperature. Cells were then permeabilized with 0.1% Triton x-100 in PBS, blocked for 1 hour with 6% BSA (Sigma, A7906) in PBS and incubated overnight at 4°C in primary antibody – mouse anti γ-H2AX 1:1000 (Genetex, GTX127342). The following day samples were allowed to reach room temperature (30-40 minutes), washed, and incubated with secondary antibody – Donkey anti-mouse AlexaFluor 488 (Abcam, ab150105) 1:2000. This was followed by washing, counterstaining with DAPI, washing, and mounting with VectaShield mounting media (Vector Laboratories, H-1000-10), and sealing with Eukitt (Sigma, 03989).

For each culture random regions were imaged at 400x magnification on a fully motorized IX-83 Olympus microscope with a Lumencor Spectra X fluorescent light system (Lumencor, Beaverton, OR, USA). The quantification of the γ-H2AX coverage of the nucleus was carried out using FIJI package (NIH, ImageJ). Briefly, all γ-H2AX channel images were brought to the same stack and the optimal threshold was determined using stack histogram and “Li” thresholding method. Then single multi-channel images were analyzed by auto-thresholding the DAPI channel to create a binary map and analyze-particle function was used to identify nuclei. The identified particle area was then used on a thresholded γ-H2AX channel to calculate the percent coverage for each cell. In total, the number of cells analyzed (for DMSO, and Etoposide, respectively): WT: 1789, 2384; *blm^Δ11/Δ11^*: 1374, 1802; *atm^Δ4/Δ4^*: 384, 414. Statistics were calculated using the Fisher exact test within genotype and FDR correction.

##### Micronuclei scoring

Cells from *blm^Δ11/Δ11^*, *atm^Δ4/Δ4^* and WT cultures were scored for micronuclei (MN). Cells were seeded at 40k cells per well in a 12-well plate with coverslips, and allowed to grow overnight. Cells were then fixed with 4% PFA in PBS (15 minutes at room temperature), washed with PBS, and stained with DAPI (1mg/ml for 15 minutes). Coverslips were mounted with VectaShield (Vector Laboratories, H-1000-10) and sealed with Eukitt (Sigma, 03989). Cells were then imaged on a Nikon Ti2E fluorescent microscope with Yokogawa W1 spinning disk, and scored manually for MN in FIJI software package for ImageJ. Specifically, DAPI positive speckles were scored as MN if: (1) they had an oval to round shape, (2) defined borders that do not overlap the main nucleus, (3) are of similar intensity as the nucleus (within 20%, above or below the nucleus intensity), (4) are between 1/3 and ∼1/16 of the main nucleus area, and (5) are not refractile or auto-fluorescent. Followed by counting the number of cells via auto thresholding and “analyze particle” function. In total, 5242 WT, 907 *blm^Δ11/Δ11^*, 3481 *atm^Δ4/Δ4^*, 2464 *cgas^Δ10/Δ10^* and 4448 *atm^Δ4/Δ^;cgas^Δ10/Δ10^*cells were scored. Data from *atm^Δ4/Δ4^* in one experiment was omitted due to over-confluency of the cells. Statistics and graphing were carried out with R programming language (4.1.3). Significance was calculated using χ2 test using the WT as expected values and FDR correction.

##### Anti-cGAMP ELISA

Cells from *cgas^Δ10/Δ10^* and WT controls were seeded at 3x10^6^ in T175 flasks. Upon reaching 80% confluency they were irradiated with 5 Gy. 48 hours after irradiation cells were trypsinized and scraped. Cells were then lyzed in Cell Extraction buffer (Invitrogen CAT#FNN001). Sample concentrations were adjusted to 4mg/ml protein using protein quantification measurements (Pierce™ BCA Protein Assay Kit, ThermoScientific, CAT#23225), following the manufacturer’s instructions. cGAMP concentration was detected with ELISA “2’,3’-Cyclic GAMP Competitive ELISA Kit” (Thermo-Fisher, CAT# EIAGAMP), and following the manufacturer’s protocol. Fluorescence was read with an Agilent Synergy H1 plate reader at a wavelength of 450 nm. Statistics and graphing were carried out with R programming language (4.1.3). Significance was calculated using one-way ANOVA with Turkey post hoc

#### Melanoma engraftment

Melanoma engraftment was performed according to^43^. Briefly, the tail fin from an 8-month-old male with a melanocyte expansion was dissected, finely minced and digested with 0.2% Collagenase Type P (Merck Millipore) and 0.12% Dispase II (Sigma Aldrich) in Leibovitz’s L-15 Medium (Gibco) for 30 min at RT. Cells were filtered using a Falcon 40 µm filter, centrifuged at 400 g for 10 minutes, resuspended in L15 with 10% FBS and injected into a *rag2* mutant fish. Following *in-vivo* expansion of the melanoma cells in the *rag2* recipient, muscle tissue containing melanoma was dissected and prepared for transplantation as described above. Fish were then monitored, and melanoma size was visually scored 5 weeks post-injection. This experiment was performed together with the experiment described in^43^ and WT controls were shared between experiments.

#### Metaphase spread preparation and FISH

For metaphase spreads, cells were treated with 0.2 ug/ml colcemid (15212-012, Gibco) for overnight, collected and incubated in pre-warmed hypotonic solution (75 mM KCl) for 7 min. The cell suspension was cytocentrifuged, washed and subsequently fixed in fixative solution (3:1 methanol: glacial acetic acid), dropped on superfrost microscope slides to prepare metaphase spreads and dried overnight before FISH. For FISH, the slides were rehydrated in PBS for 5-10 min, treated with PBS containing 4% formaldehyde for 2 min, washed 3 times with PBS, 5 min each, incubated in pepsin solution (1 mg/ml at pH 2.0 acidified water), for 10 min at 37°C, washed again with PBS, fixed in 4% formaldehyde for another 2 min, washed 3 times in PBS, 5 min each, dehydrated with 70%, 90%, and 100% ethanol (5 min each) and air-dried. Hybridization mix (70%formamide, 10 mM Tris·HCl pH 7.2, and 10% NEN blocking solution, 50mM Tris PH7.2, 8% MgCl2 buffer, 10ng telomeric PNA-(CCCTAA)3Cy3 probe (F1002, Panagene) was denatured for 5 min at 90°C, then the slides were denatured in hybridization mix for 3 min at 80°C on a heating block. After denaturation, hybridization was continued for 2h at RT in the dark. Slides were washed twice, 15 min each, in PBS containing 70% formamide, and washed 3 times with (1 M Tris PH7.4, 6 ml 5M NaCl, 100 ul Tween 20), 5 min each. Lastly, the slides were dehydrated with 70%, 90%, and 100% ethanol (5 min each), and mounted with mounting media containing DAPI (VE-H-1200, Zotal). Significance was calculated using one-way ANOVA with Tukey post hoc. Microscopy, plotting, statistics.

#### Data availability

All raw and processed RNA sequencing data can be found in the Gene Expression Omnibus (GEO) database (GSE270096). All other data are available from the corresponding author upon request.

#### Code availability

The code supporting the current study is available in the following GitHub repository https://github.com/Harel-lab/cGAS-STING-pathway-in-killifish.

